# Photo-crosslinking hERG channels causes a U.V.-driven, state-dependent disruption of kinetics and voltage dependence of activation

**DOI:** 10.1101/2024.01.09.574834

**Authors:** Sara J. Codding, Matthew C. Trudeau

**Author notes:** Correspondence to Matthew C. Trudeau.

## Abstract

Human ether-à-go-go related gene (hERG) voltage-activated potassium channels are critical for cardiac excitability. Characteristic slow closing (deactivation) in hERG is regulated by direct interaction between the N-terminal Per-Arnt-Sim (PAS) domain and the C-terminal cyclic nucleotide binding homology domain (CNBHD). We aim to understand how the PAS domain that is distal to the pore rearranges during gating to allosterically regulate the channel pore (and ion flux). To achieve this, we utilized the non-canonical amino acid 4-Benzoyl-L-phenylalanine (BZF) which is a photo-activatable cross-linkable probe, that when irradiated with ultraviolet (U.V.) light forms a double radical capable of forming covalent cross-links with C-H bond-containing groups, enabling selective and potent U.V.-driven photoinactivation of ion channel dynamics. Here we incorporate BZF directly into the hERG potassium channel PAS domain at three locations (G47, F48, and E50) using TAG codon suppression technology. hERG channels with BZF incorporated into the PAS domain (hERG-BZF) showed a significant change in the biophysical properties of the channel. hERG-G47BZF activated slowly when irradiated in the closed state (-100mV) but deactivated quickly when irradiated in both the open (0mV) and closed state. hERG-F48BZF channels showed a state independent and U.V. dose-dependent change in channel activation (slowing down) and channel deactivation (speeding up), as well as a marked change (right-shift) in the voltage-dependence of conductance. When irradiated at -100 mV hERG-E50BZF showed a state dependent and U.V. dose-dependent change in a channel activation (slowing down) and deactivation (speeding up) of channel deactivation, as well as a marked change (right-shift) in the voltage-dependence of conductance that occurred only when the channel was irradiated in the closed state (-100mV). This approach demonstrated that direct photo-crosslinking of the PAS domain in hERG channels causes a measurable change in biophysical parameters and more broadly stabilized the closed state of the channel. We propose that altered channel gating is as a direct result of reduced dynamic motions in the PAS domain of hERG due to photo-chemical crosslinking.

## Introduction

The KCNH family of potassium voltage-gated ion channels is comprised of EAG (ether-à-go-go, KCNH1), ERG (ether-à-go-go related, KCNH2), and ELK (ether-à-go-go like; KCNH3). KCNH channels share close homology to cyclic nucleotide modulated channels HCN (hyperpolarization-activated cyclic nucleotide regulated) and CNG (cyclic nucleotide-gated) channels rather than to other voltage-gated potassium channels (Fig.1)(1). Structurally, KCNH channels have six transmembrane segments (Fig.1): S1-S4 form the voltage sensing domain (VSD) (Fig.1-grey, purple), S5-S6 and pore loop (P) form the ion conduction pathway (Fig.1-black). Intracellularly, KCNH channels have a N-terminal Per-Arnt-Sim (PAS) domain (Fig.1-green), an N-linker (Fig.1-pink) that connects the PAS to the VSD, and a C-terminal cyclic nucleotide binding homology domain (CNBHD) (Fig. 1 blue), within which a small intrinsic ligand (IL) motif is nested (Fig.1-orange), and a C-terminus (Fig.1-brown)(2). Four subunits assemble to make a tetrameric channel. KCNH channels share similar domain architecture to HCN and CNG channels, yet unlike HCN and CNG, they are not directly activated by cyclic nucleotides (3, 4). Instead, in KCNH channels the IL motif resides within a pocket of the CNBHD that is homologous to the nucleotide biding site of HCN/CNG channels (Fig.1)(5).

**Figure 1:**
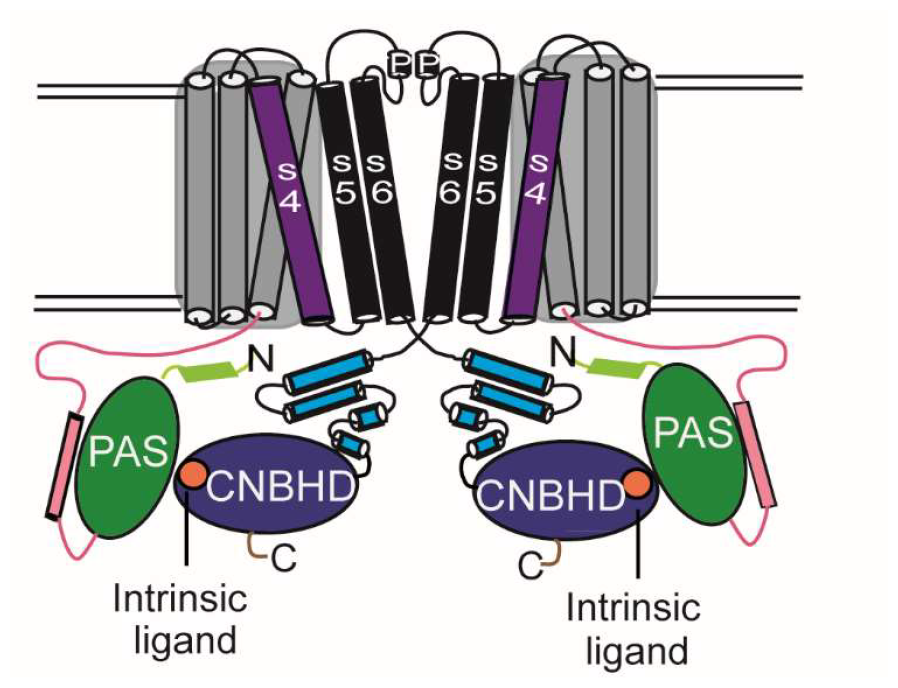
Cartoon of the hERG potassium ion channel, only two subunits included for clarity. Labeled from N terminus to C terminus: In light green shows the N terminal Per-Arnt-Sims (PAS)-Cap domain; Green-PAS domain, Pink-N-linker; Grey-voltage sensor domain helices S1-S3 and Purple-charge sensing S4 helix; Black-Pore domain composed of S5 helix, followed the pore loop (P), and S6 helix; Cyan-N-linker; Blue-cyclic nucleotide biding domain (CNBHD); Orange-intrinsic ligand; Brown-C-terminus.

The KCNH channel human ether-à-go-go-related gene (hERG, KCNH2) is critical for cardiac function and encodes the alpha subunit of the delayed-rectifier (IKr) current that drives action potential repolarization in the heart (6-8). Inherited mutations in human ERG (hERG) cause familial Type 2 Long QT syndrome (LQT2), a predisposition to cardiac arrhythmias and sudden death (6, 9, 10). The primary mechanism for acquired LQT syndrome is inhibition of hERG from off-target effects of pharmaceuticals and this is a common clinical problem (11, 12).

In hERG, interactions between the N-terminal Per-Arnt-Sim (PAS) of one subunit and the C-terminal Cyclic Nucleotide Binding Homology Domain (CNBHD) of a neighboring subunit has been implicated in the characteristic slow closing of the channel gate (13, 14). Yet, how domains distal to the pore rearrange during gating to regulate channel the pore (and ion flux) is unclear. Historically, work to study the electromechanical couplings in voltage gated ion channels (VGIC) utilized high affinity metal bridges which require dual cysteine mutations within favorable structural proximity that when exposed to cadmium, form a metal bridge, thereby locking a channel in a particular conformational state (15). This approach is limited as it requires several permissive cysteine mutations, cytosolic placement of cysteines for metal accessibility, and detailed structural information for successful interpretation of results.

Here, we utilize a highly effective approach to photochemically crosslink hERG channels using an ultraviolet (U.V.) reactive non-canonical amino acid (ncAA) benzoyl-L-phenylalanine (BZF) (Fig.2), that can be incorporated into growing nascent peptides with TAG codon suppression technology (Fig.3) (16, 17). A pulse of U.V. light applied to BZF transforms it into a reversible double radical capable of forming a new carbon-carbon bond to neighboring atoms. When this is performed in tandem with electrophysiology to hold the channel in various conductive (and conformational) states, BZF can capture/lock conformational states of domains within the channel. Examining the gating effects due to crosslinking can then inform on allosteric of effects of domains distal to the pore gate on channel gating and voltage sensitivity. This approach is advantageous as BZF can i) be incorporated into proteins at locations inaccessible to metal bridging ii) provide information on interactions between regions of proteins and if they occur in a state dependent manner iii) BZF provides information on the distribution of channel conformational states under distinct voltage clamp potentials, and this is encoded within the rate and efficacy of UV crosslinking(18).

**Figure 2:**
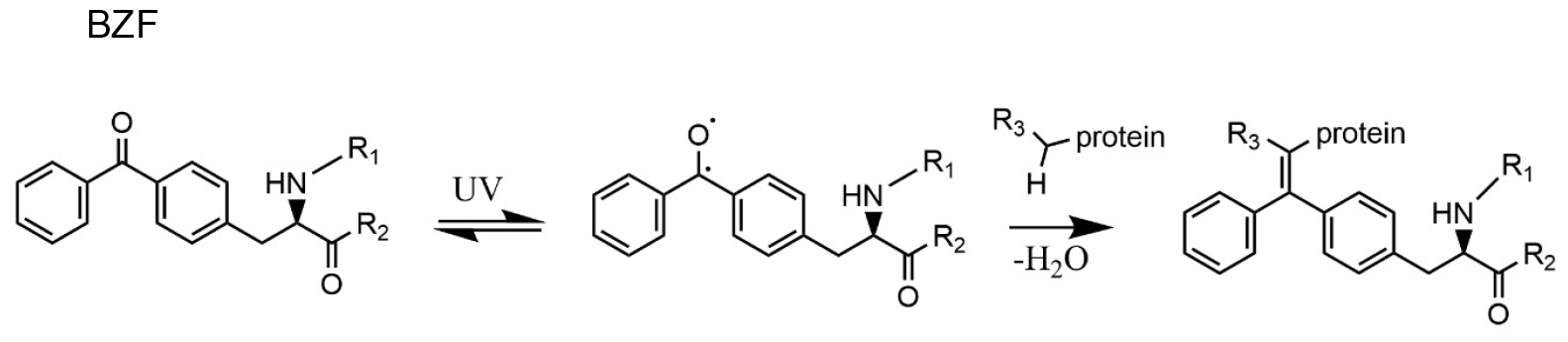
BZF double radical formation. BZF (left) when incorporated into a protein will be part of the primary peptide chain with the amino group forming a peptide bond and bound to the to the upstream endogenous amino acid (R_1_) and the carboxyl group bound to the endogenous downstream amino acid (R_2_), and harbors the functional group (benzophenone) attached to Cα. When exposed to UV light, BZF forms double radical capable of a.) relaxing to the ground state or b.) eliminating water and forming a new carbon-carbon bond depicted here with a protein, where R_3_ is the moiety disrupted by new bond formation and is dependent on localized chemistry.

**Figure 3:**
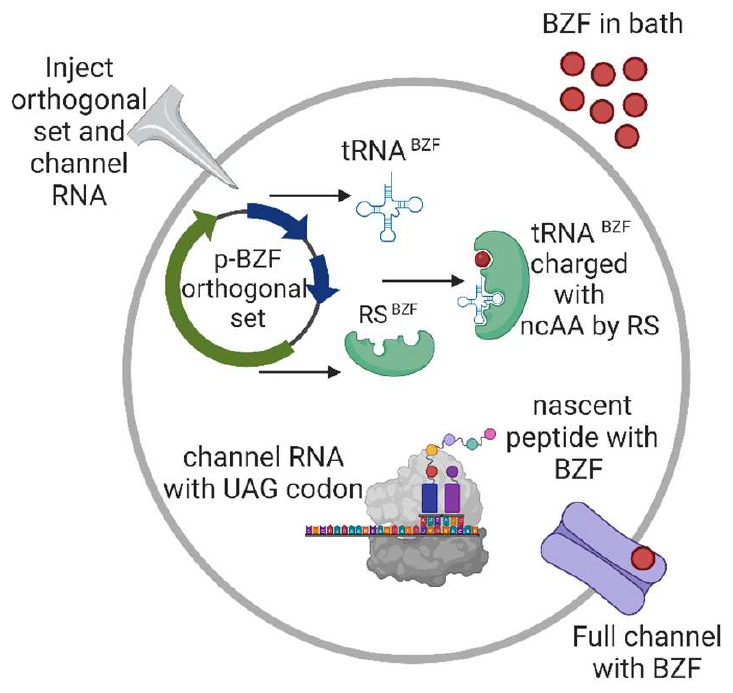
Diagram of BZF incorporation. Oocyte depicted by large circle that serves as orthogonal expression system. Oocyte is injected (blind nuclear) with the plasmid that contains the genetic information for the transfer RNA (tRNA^BZF^ schematic) and RNA synthetase (RS^BZF^ green), also known as the orthogonal set, bind and then charge tRNA with BZF. The charged tRNA^BZF^ is used by the ribosome (gray cartoon) to read through a genetically modified version of hERG RNA containing a “TAG” stop (or amber stop) codon and insert BZF into the nascent peptide is translocated to the membrane as a full channel (purple) with BZF incorporated (red circle).

Here, we report that BZF is successfully incorporated into the PAS domain without a large perturbation of channel function. Photo crosslinking hERG-BZF channels we report i) a state independent (hERG-G47BZF, hERG-F48BZF) and ii) state dependent change (hERG-G47BZF, hERG-E50BZF) of the voltage dependence of conductance, channel kinetics, and/or both, measured with inside out patch clamp electrophysiology. With photo crosslinking we show that the conformation of the PAS domain in hERG channels allosterically regulates the channel gate.

## Results

### Screening hERG-TAG channels for incorporation of BZF into the PAS domain

For this study we expressed human ERG channel Xenopus oocytes. To test if dynamics of the PAS domain allosterically regulates hERG channel gating, we chose to incorporate the photochemical crosslinking ncAA BZF directly into the PAS domain of hERG channels (Fig.4, A). To achieve this, we placed TAG codons in the hERG gene at sites of interest within the PAS domain corresponding to the amino acids G47, F48, and E50 in a small solvent exposed helix spanning residues 46-51 (Fig.4 A,B). These residues were chosen with the rationale that they were within structural proximity to the i) CNBHD domain to potentially crosslink between subunits *and/or* ii) have rotamers that could crosslink within the PAS domain itself.

**Figure 4:**
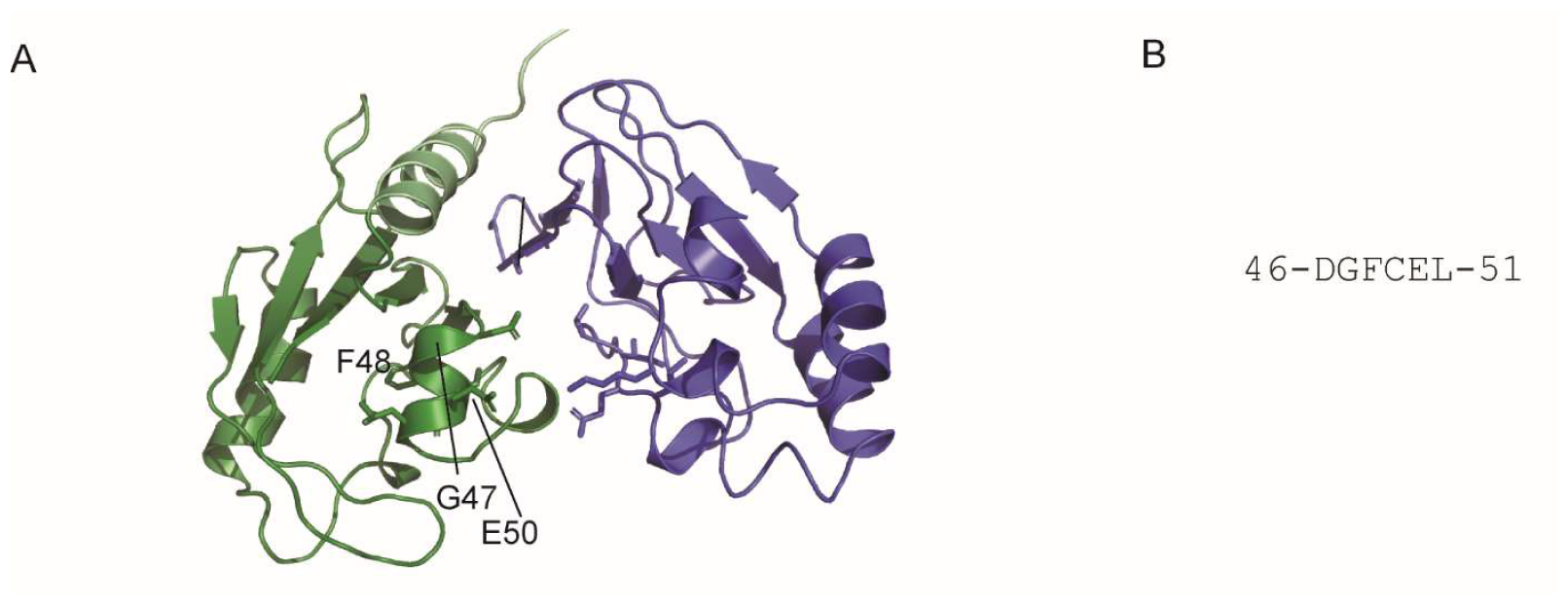
Placement of BZF in hERG channels. A) Structure of hERG channel PDB 5VA2 depicting intracellular domains of neighboring subunits, PAS (Green) and CNBHD (Blue). Locations of residues in the PAS domain helix spanning residues 46 to 50 for incorporation of BZF indicated by labeling, G47, F48, and E50 and the side chains shown as sticks where applicable. B) Primary sequence of PAS helix spanning residues 46-50.

To incorporate BZF into hERG channels for cross linking studies we utilized TAG codon suppression technology that utilizes the least common stop codon (TAG or Amber stop) (Shultz 2007, Liu brock) (Fig.3). In brief, to utilize this technology for heterologous expression in oocytes, a TAG codon is placed into the hERG gene at the location of interest for BZF incorporation. The gene is converted to RNA via reverse transcriptase and injected into the cytosol of the oocyte. Orthogonal machinery encoding a tRNA/synthetase set are blind injected into nucleus of the oocyte. The oocyte is incubated with BZF in the media. The orthogonal tRNA synthetase recognizes and attaches BZF to the orthogonal tRNA, and the BZF is incorporated by the ribosome by suppression of the TAG codon in hERG channel mRNA and make full length channel with BZF incorporated into the site of interest (Fig.3).

To rule out any nonspecific protein expression with the hERG-TAG constructs such as: i) non-specific read through by skipping the TAG codon ii) nonspecific charging of orthogonal or endogenous-tRNAs by the orthogonal or endogenous tRNA synthetases, we expressed our channels in the presence of the orthogonal set and the absence (+/-) of the ncAA BZF. In addition, we expressed channels in the presence of the orthogonal set and in the presence of ncAA BZF (+/+). We then screened for expression and similar biophysical properties of hERG and hERG-TAG channels under these conditions 3 days post injection (3dpi) with two-electrode voltage-clamp.

We observed no change in expression of hERG channels under both (+/-) and (+/+) conditions (Fig.5). For hERG-TAG constructs, we observed no meaningful expression in (+/-) conditions, and robust expression in (+/+) conditions (Fig.5) 3dpi.

**Figure 5:**
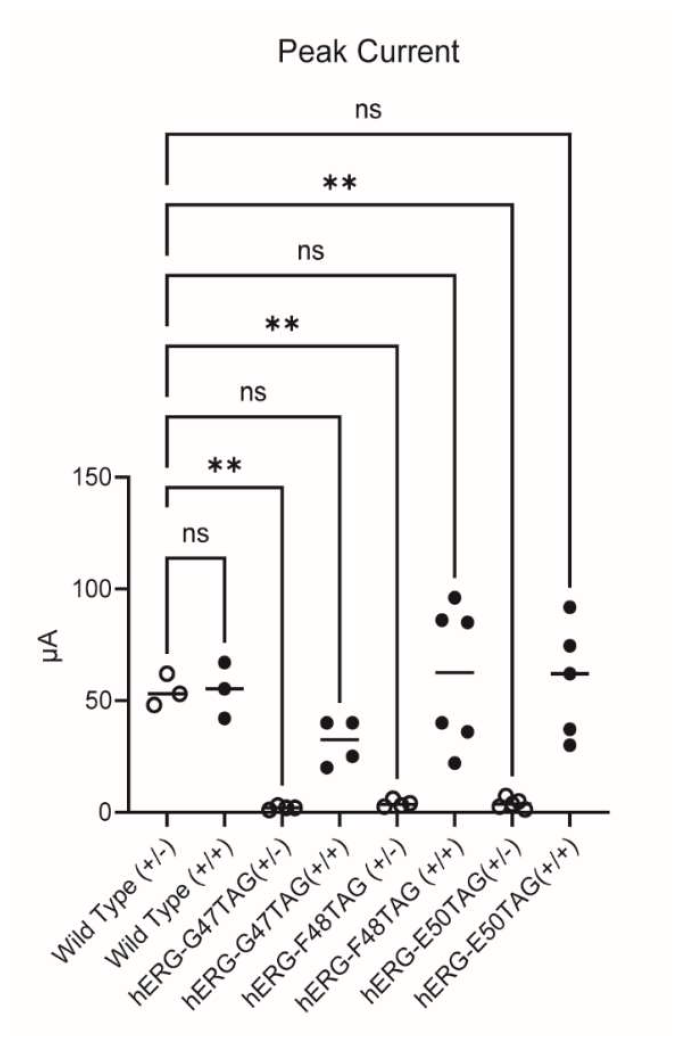
Peak Current Measured with two-electrode voltage clamp (TEVC) hERG constructs harboring a TAG codon were screened for specificity of TAG codon suppression. Oocytes with injected hERG channels were also subjected to nuclear injection of the orthogonal set and allowed to incubate for 3 days post injection (3DPI) in the presence (+/+) (black filled circles) or absence (+/-) (black hollow circles) BZF in the bath solution. A family of traces was recorded with TEVC and peak current reported at 40mV potential.

### hERG channel currents are U.V. insensitive

To test if hERG channels were photo sensitive to U.V. light, excised membrane patches from oocytes expressing hERG in the presence of the orthogonal set and BZF were subjected to U.V. irradiation and the biophysical properties were recorded using inside out patch clamp electrophysiology.

Specifically, excised patches were clamped at a known holding voltage (-100mV or 0mV) and hERG currents were recorded (0sec U.V.), then exposed to short discrete (and increasing in duration) pulses of U.V. irradiation for total exposure time intervals of 1sec, 3sec, 7sec, 15sec, and channel currents were recorded between each U.V. pulse.

hERG channels showed characteristic slow activation and deactivation in excised inside out patches when a family of current traces was recorded when irradiated at a -100mV holding potential (Fig.6, A). There was no observable change in hERG channel currents after 15 seconds of U.V. irradiation under these conditions (Fig.6,B). Additionally, hERG channels showed characteristic slow activation and deactivation kinetics in excised inside-out patches when a family of current traces was recorded when irradiated at 0mV holding potential (Fig.6, C). There was no observable change in hERG channel currents after 15 seconds of U.V. exposure under these conditions (Fig.6,D). The voltage protocol was indicated in (Fig.6, E).

The conductance-voltage relationship was plotted for all currents recorded at the discrete U.V. irradiation intervals. Regardless of U.V. exposure duration, or the holding potential that the channel was clamped prior to irradiation, we observed no change the G-V curve (Fig. 6, F,G). The change in the 50% conductance voltage for each condition was plotted and no statistically significant difference in the ΔV0.5 was observed (Fig.6, H).

**Figure 6:**
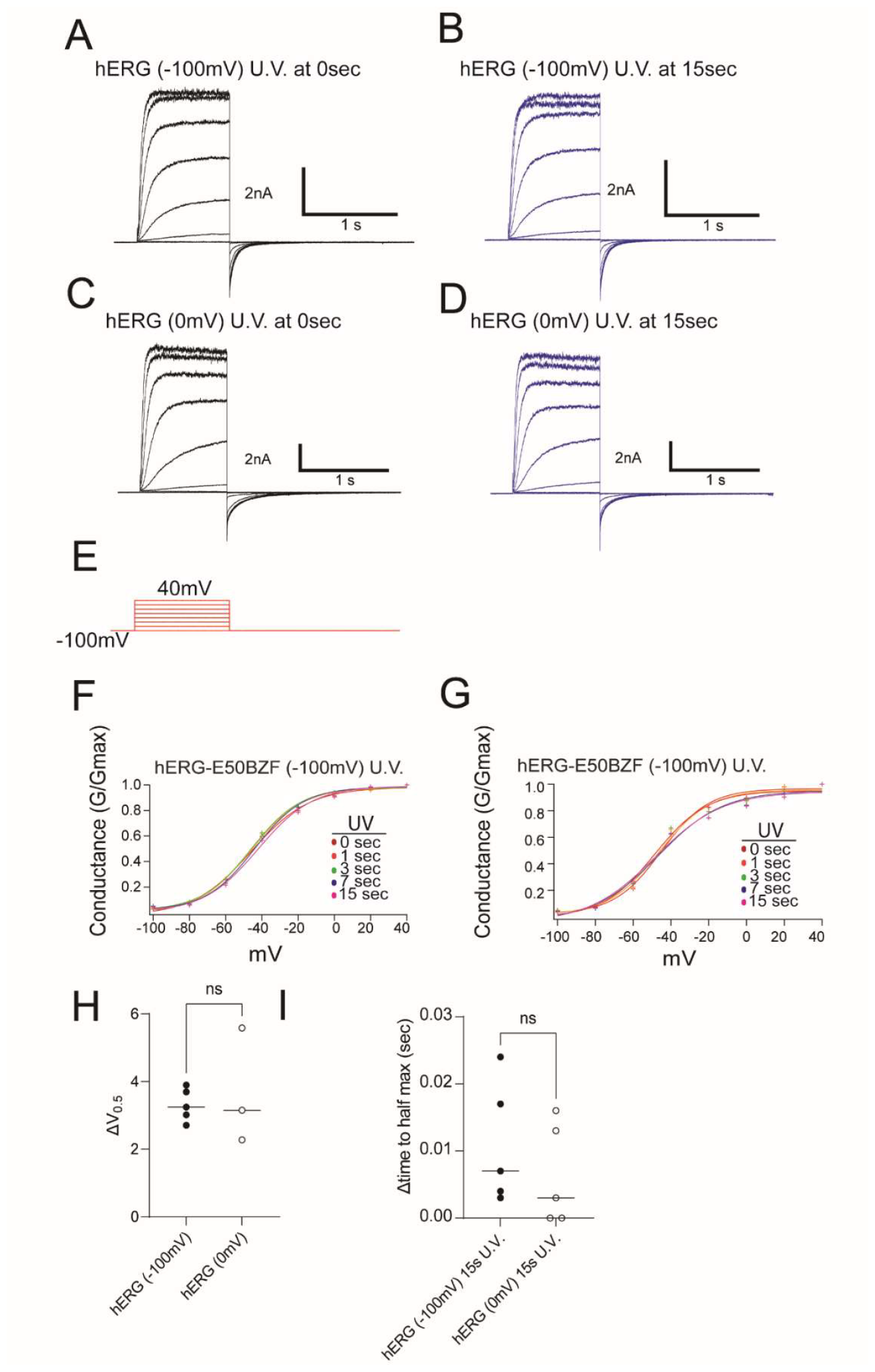
hERG channel biophysical properties of conductance and activation before and after U.V. irradiation. A.) Current family of traces recorded from inside out excised patches containing hERG channels held at -100mV prior, and during, U.V. exposure (U.V. 0 sec). B.) Family of traces as in panel A after 15sec cumulative exposure of U.V. irradiation (blue). C.) Family of traces as in panel A with channels held open (0mV) and prior to U.V. irradiation D.) Family of traces as in panel C with channels held open (0mV) and after 15sec cumulative exposure to U.V. irradiation. E.) Voltage protocol for A-D F.) Conductance Voltage curves plotted before (0 seconds) and after 1 second, 3 seconds, 8 seconds, 15 seconds exposure to U.V. irradiation and no dose dependent change is observed with the channels held at -100mV during U.V. irradiation. G.) Conductance Voltage curves plotted before (0 seconds) and after 1 second, 3 seconds, 8 seconds, 15 seconds exposure to U.V. irradiation and no dose dependent change is observed with the channels held at 0mV during U.V. irradiation. H.) Change in V0.5 after 15 seconds U.V. exposure as a function of holding potential during U.V. irradiation (-100mV (filled circles), 0mV (empty circles)). I.) Change in time to apparent half maximum current of voltage trace (-40m). opening potentials, (black: -100mV irradiation, blue: 0mV irradiation)

Channel activation, or the apparent time to half maximum voltage for the channel opening at voltage step to -40mV, showed no change as a function of U.V irradiation regardless of holding potential of the channel during irradiation (-100mV or 0mV) (Fig.6, I). This was also true for channel deactivation, where tau1 showed no change after U.V. irradiation regardless of the voltage holding potential during U.V. exposure (Fig.7, A-D, G).

**Figure 7:**
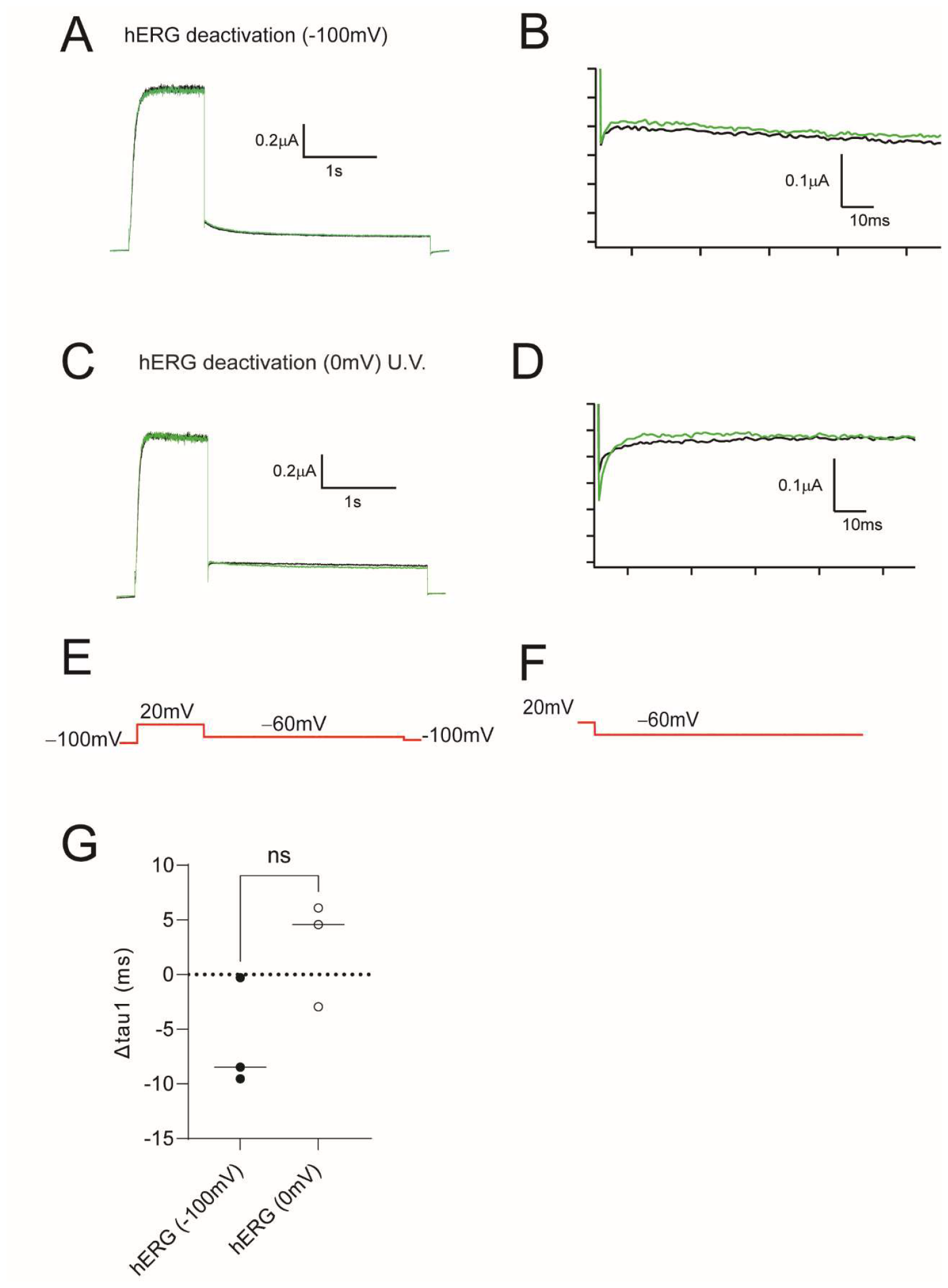
hERG channel biophysical properties of deactivation before and after U.V. irradiation. A.) Currents recorded from inside out excised patches containing hERG channels to measure rate of channel closure at -60mV. Channels held at -100mV, before U.V. exposure (black) and after 15 seconds U.V. irradiation (green), normalized. B.) The first 60ms of trace (zoomed in for clarity) of currents in panel A after channel closure at -60mV C.) Currents recorded from inside out excised patches containing hERG channels to measure rate of channel closure at -60mV. Channels held at 0mV, before U.V. exposure (black) and after 15 seconds U.V. irradiation (green), normalized. D.) The first 60ms of trace (zoomed in for clarity) after channel closure at -60mV of currents in panel C. E. Voltage protocol for panel A, C. F.) Voltage protocol for panels B, D. G.) Change in tau1 of deactivation of hERG channels after U.V. irradiation at respective irradiation holding potentials.

To investigate the underlying role of PAS domain dynamics in hERG gating, we performed excised patch clamp recordings of hERG-BZF channels with BZF incorporated at G47, F48 and E50 of the PAS domain. We compared changes in the biophysical properties of hERG-BZF channels to hERG channels without BZF. For all hERG-BZF channels, excised membrane patches from oocytes expressing hERG-BZF channels in the presence of the orthogonal set and BZF were subjected to U.V. irradiation and the biophysical properties were recorded using inside out patch clamp electrophysiology as noted for hERG channels above.

### hERG-G47BZF photochemical crosslinking slows channel activation in a U.V dependent and state dependent manner, and speeds channel deactivation in a U.V. dependent and state independent manner

We measured hERG-G47BZF channels before and after U.V irradiation. We observed slow activation and deactivation in hERG-G47BZF channels before photochemical crosslinking (Fig.10, A,C). However, after exposure to U.V. irradiation in the closed state (-100mV), hERG-G47BZF channels were slower to activate at low voltages (-40mV) (Fig.10, B), but this effect was not observed when hERG-G47BZF channels were irradiated in the open state (0mV) (Fig.10, I).

Additionally, there was no measurable change in the conductance-voltage relationship regardless of the state of the channel during U.V. irradiation (Fig.10, F-H). Comparing the ΔV0.5 of hERG-G47BZF to hERG channels we observed no statistical difference (Fig.10, H).

hERG-G47BZF channels deactivated faster after 15 sec of U.V. irradiation regardless of the state of the channel during U.V. irradiation (-100mV or 0mV) (Fig.11, A-D). The change in hERG-G47BZF channel deactivation rate compared to hERG channels was significant and state independent (Fig.11, G).

### hERG-F48BZF photochemical crosslinking slows channel activation, speeds deactivation, and right shifts the voltage dependence of activation in a U.V dependent and state independent manner

We recorded from hERG-F48BZF and after U.V. irradiation, the biophysical properties of hERG-F48BZF channels were altered (Fig.12, A-D). We measured a marked right shift of the voltage dependence of conductance that was U.V. dependent and state independent (Fig.10, F,G). The change in the ΔV0.5 of hERG-F48BZF was significant compared to hERG channels (Fig.10,H).

Additionally, the hERG-F48BZF channel, similar to hERG-G47BZF channels, were slower to activate at low voltages (-40mV) (Fig.10, B,D). The change in apparent time to half maximum voltage at -40mV after 15 seconds of U.V. irradiation for hERG-F48BZF channels was state independent and significantly larger than that observed for hERG channels. But unlike hERG-G47BZF, in hERG-F48BZF channels this effect was observed regardless of the state in which the channel was irradiated (-100mV, or 0mV) (Fig.10, I).

hERG-F48BZF channels also exhibited a large change in channel deactivation after U.V. irradiation (Fig.11, A-D,G). hERG-F48BZF deactivated faster after U.V. irradiation regardless of holding voltage during U.V. exposure. Compared to hERG channels, the change in hERG-F48BZF channel deactivation rate (faster, speeding up of tau1) after U.V. irradiation was significant, state independent, and observed when the channel was irradiated at both -100mV and 0mV.

### hERG-E50BZF photochemical crosslinking slows channel activation, speeds deactivation, and right shifts the voltage dependence of activation in a U.V dependent and state dependent manner

We recorded from hERG-E50BZF channels and irradiated the channels as noted above, at discrete intervals, at either -100mV or 0mV holding potentials. Compared to hERG channels the change in biophysical properties of hERG-E50BZF channels occurred both in the steady-state properties and kinetics properties and was state dependent.

hERG-E50BZF channel currents prior to U.V. irradiation (Fig.12) compared to after 15 seconds of total U.V. irradiation at -100mV holding potential (closed state), show a slowed activation at low voltages (-40mV) and faster channel deactivation (Fig.12, B). This effect was not observed for hERG-E50BZF channels irradiated at 0mV for 15 seconds (Fig.12,D).

The conductance-voltage relationship of hERG-E50BZF was plotted for the family of current traces collected after U.V. irradiation intervals (Fig.12 F,E). For hERG-E50BZF channels irradiated at -100mV (closed state) showed a U.V. dose dependent right shift of the voltage dependence of conductance of the channel. This effect was not observed when hERG-E50BZF channels were irradiated at 0mV (open state). Compared to hERG, the change in the 50% conductance voltage for hERG-E50BZF was significant when the irradiated in the closed state but not the open state of the channel.

We also observed a state dependent change in the time to apparent half maximum voltage of hERG-E50BZF channels (closed state), where the channels open slower at low voltages (-40mV) when irradiated in at -100mV.

hERG-E50BZF channels exhibited a large change in channel deactivation after U.V. irradiation at -100mV, but not 0mV (Fig.13, A-D,G). hERG-E50BZF deactivated faster after U.V. irradiation at -100mV. We observed no change in deactivation at 0mV holding voltage during U.V. exposure. Compared to hERG channels, the change in hERG-E50BZF channel deactivation rate (faster, speeding up of tau1) after U.V. irradiation at -100mV was significant, state dependent (closed state), and observed when the channel was irradiated at -100mV, but not at 0mV.

## Discussion

Here we show that i) hERG channels are U.V. insensitive ii) hERG channels with Amber mutations in the PAS domain (Fig.4) were rescued by the photoactivatable ncAA BZF (Fig.5) and iii) photo crosslinking hERG-BZF channels stabilized the closed state of the channel, to differential extents depending on BZF location, in a U.V. dependent manner, as measured with excised patch clamp electrophysiology.

hERG channels expressed in the presence of BZF and the orthogonal set (Fig.3), then irradiated with U.V. light did not exhibit a measurable photo reactive change in channel biophysical properties with patch clamp electrophysiology. Neither the steady-state voltage dependence of conductance, nor the kinetics of activation or deactivation were markedly changed after exposure to U.V. light (Fig.6,7). This result ruled out measurable i) nonspecific BZF incorporation into hERG and ii) photosensitivity of hERG channels to U.V. light. This suggests that BZF rescue of Amber mutant channels is specific and the observed changes after U.V. irradiation is associated with formation of photochemical crosslinks of hERG-BZF channels. Thus, we were able to compare changes in steady-state and kinetic properties of hERG-BZF channels to hERG and explicitly attribute those changes to photochemical crosslinking (19).

Our findings demonstrated that in hERG-G47BZF channels, photochemical crosslinking slowed channel activation in a U.V dependent and state dependent manner and increased the rate of channel deactivation in a U.V. dependent and state independent manner (Fig.8,9). This result suggests that motions in the PAS domain that facilitate channel activation are inhibited by photochemical crosslinking hERG-G47BZF channels in the closed state. But, that this result is not observed at higher voltages suggests that the effect is overcome by a larger voltage driving force. Additionally photochemical crosslinking hERG-G47BZF channels appears to have the same effect on tau1 regardless of the holding potential during crosslinking as the change in tau1 is observed to occur in state independent manner. This suggests that motions in the PAS domain that regulate activation are not identical to those that regulate deactivation, which follows the paradigm that the pathway to channel deactivation isn’t the reverse of channel activation (20, 21).

**Figure 8:**
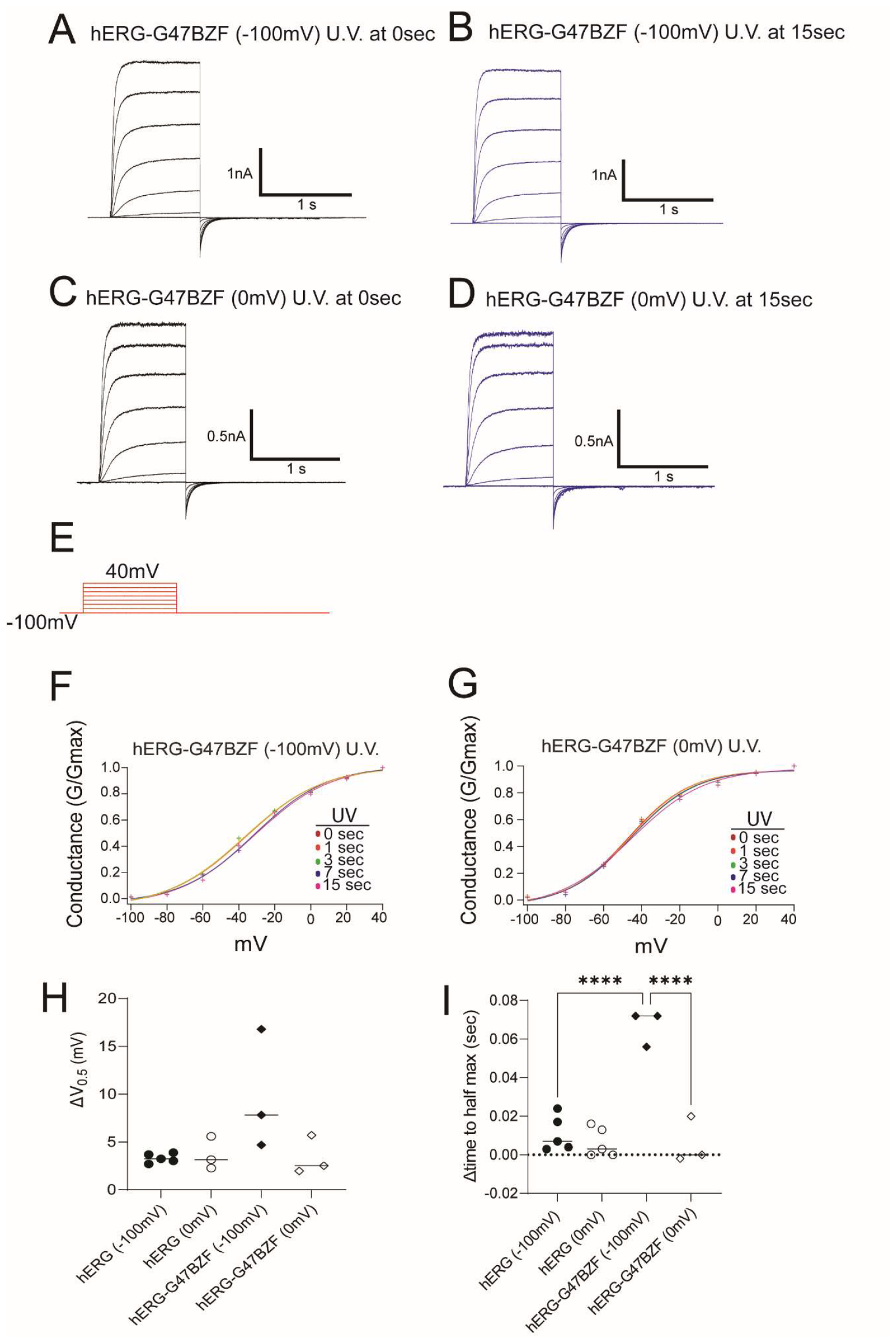
hERG-G47BZF channel biophysical properties of conductance and activation before and after U.V. irradiation. A.) Current family of traces recorded from inside out excised patches containing hERG-G47BZF channels held at -100mV prior, and during, U.V. exposure (U.V. 0 sec). B.) Family of traces as in panel A after 15sec cumulative exposure of U.V. irradiation. C.) Family of traces as in panel A with channels held open (0mV) and prior to U.V. irradiation D.) Family of traces as in panel C with channels held open (0mV) and after 15sec cumulative exposure to U.V. irradiation. E.) Voltage protocol for A-D. F.) Conductance Voltage curves plotted before (0 seconds) and after 1 second, 3 seconds, 8 seconds, 15 seconds exposure to U.V. irradiation and no dose dependent change is observed with the channels held at -100mV during U.V. irradiation. G.) Conductance:Voltage curves plotted before (0 seconds) and after 1 second, 3 seconds, 8 seconds, 15 seconds exposure to U.V. irradiation and no dose dependent change is observed with the channels held at 0mV during U.V. irradiation. H.) Change in V0.5 after 15 seconds U.V. exposure (filled symbols:-100mV, empty symbols: 0mV holding potential). I.) Time to apparent half maximum current of voltage trace (-40mV opening potentials, (black filled: -100mV irradiation, black empty: 0mV irradiation).

**Figure 9:**
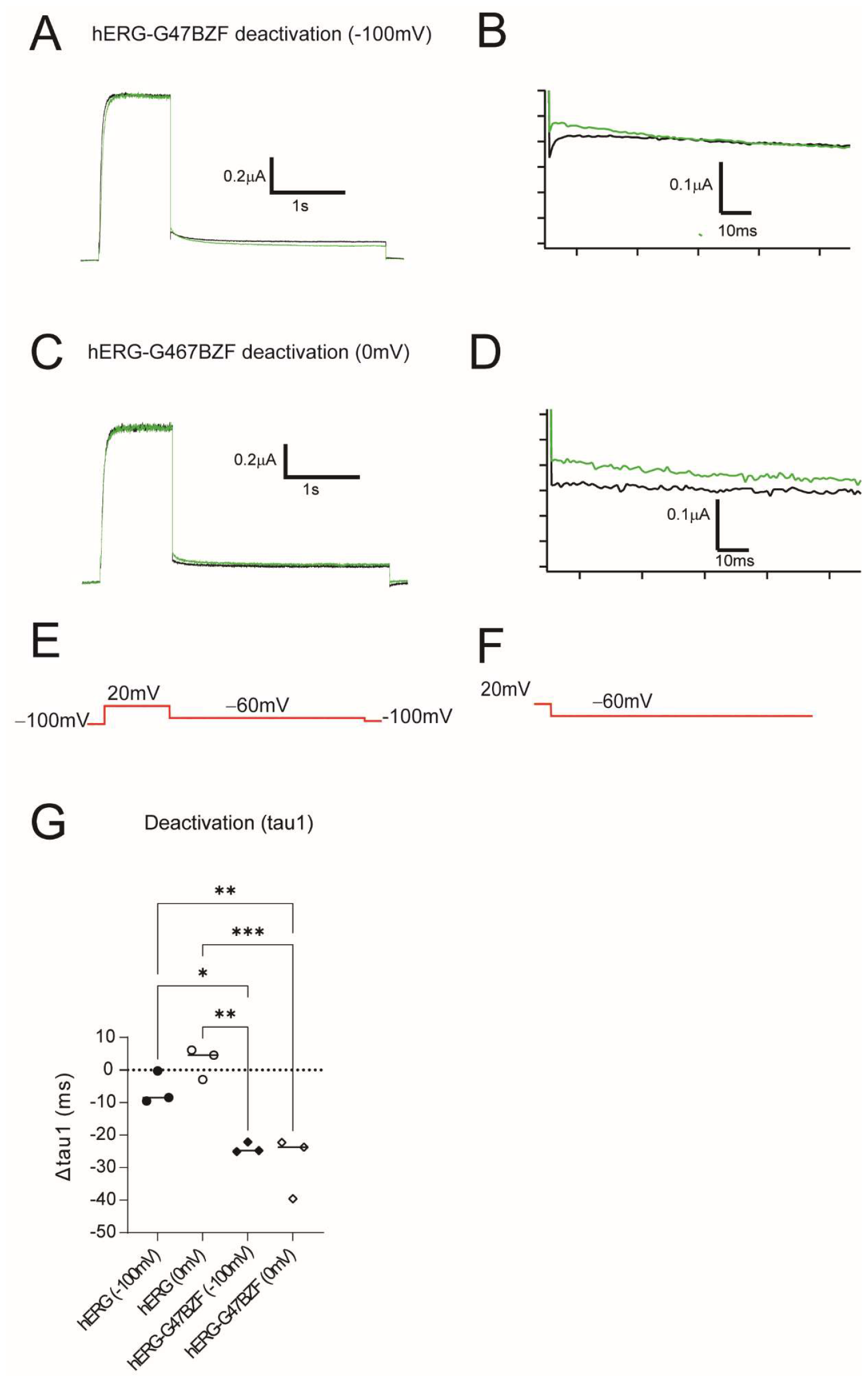
hERG-G47BZF channel biophysical properties of deactivation before and after U.V. irradiation. A.) Currents recorded from inside out excised patches containing hERG-G47 channels to measure rate of channel closure at -60mV. Channels held at -100mV, before U.V. exposure (black) and after 15 seconds U.V. irradiation (green), normalized. B.) The first 60ms of trace (zoomed in for clarity) of currents in panel A after channel closure at -60mV C.) Currents recorded from inside out excised patches containing hERG channels to measure rate of channel closure at -60mV. Channels held at 0mV, before U.V. exposure (black) and after 15 seconds U.V. irradiation (green), normalized. D.) The first 60ms of trace (zoomed in for clarity) after channel closure at -60mV of currents in panel C. E. Voltage protocol for panel A, C. F.) Voltage protocol for panels B, D. G.) Tau1 of deactivation of hERG-G47BZF channels before (black-filled) and after (black-empty) U.V. irradiation at respective holding potentials.

In hERG-F48BZF, photochemical crosslinking also slowed channel activation, increased the rate of deactivation, and right shifted the voltage dependence of activation in a U.V dependent and state independent manner (Fig. 10,11). This suggested that the PAS domain motions altered by crosslinking hERG-F48BZF are state independent and play a role in channel activation at low voltages. Further, hERG-F48BZF motions altered by crosslinking affect the voltage sensitivity of the channel observed by a marked right shift of the voltage dependence of conductance. hERG-F48BZF crosslinking destabilized the open state of the channel as we observed a state independent change and increase in channel deactivation rate. Finally, loss of PAS motions in hERG-F48BZF altered both steady-state and kinetic properties of the channel.

**Figure 10:**
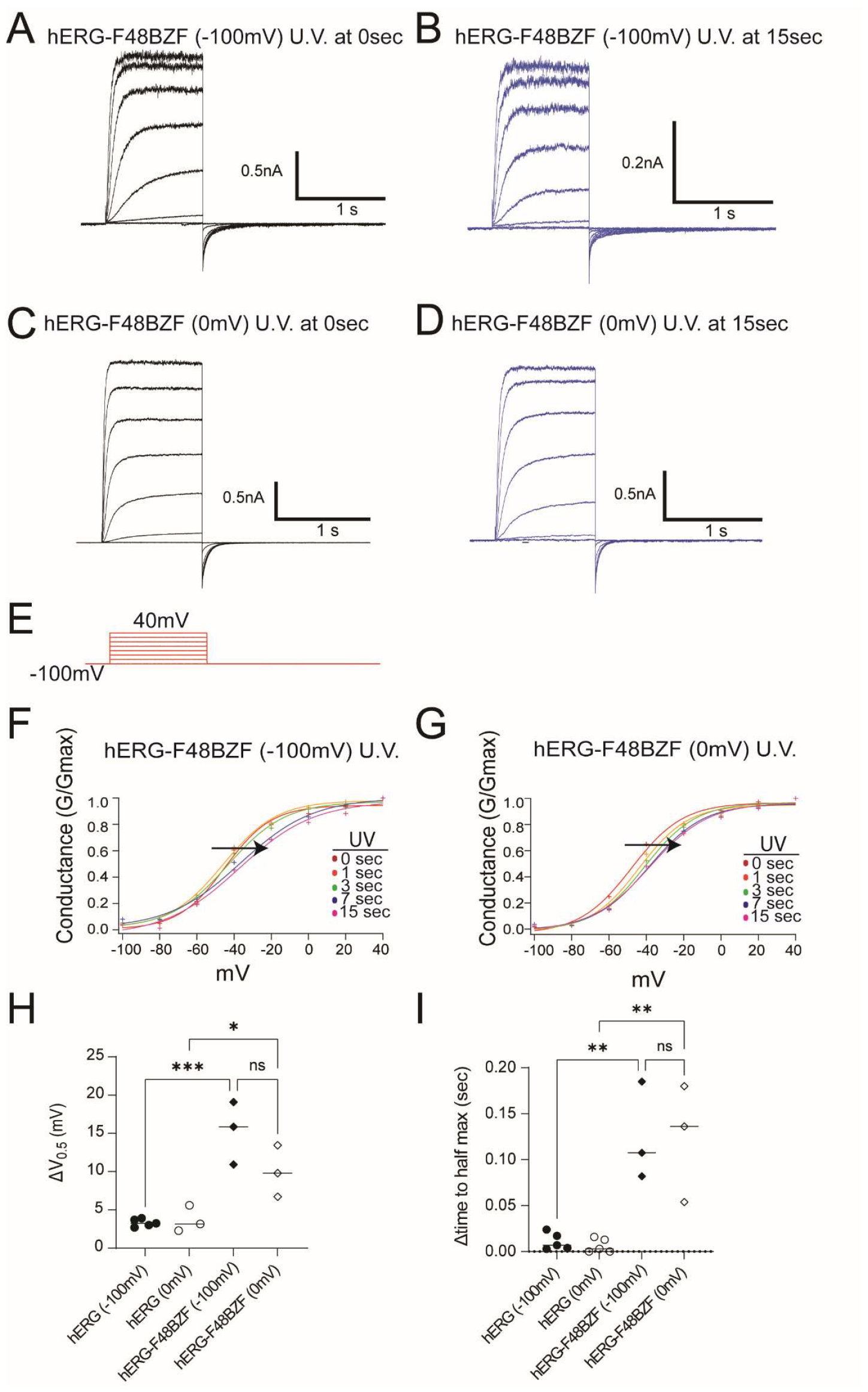
hERG-F48BZF channel biophysical properties of conductance and activation before and after U.V. irradiation. A.) Current family of traces recorded from inside out excised patches containing hERG-F48BZF channels held at -100mV prior, and during, U.V. exposure (U.V. 0 sec). B.) Family of traces as in panel A after 15sec cumulative exposure of U.V. irradiation. C.) Family of traces as in panel A with channels held open (0mV) and prior to U.V. irradiation D.) Family of traces as in panel C with channels held open (0mV) and after 15sec cumulative exposure to U.V. irradiation. E.) Voltage protocol for A-D. F.) Conductance Voltage curves plotted before (0 seconds) and after 1 second, 3 seconds, 8 seconds, 15 seconds exposure to U.V. irradiation and a dose dependent change is observed with the channels held at -100mV during U.V. irradiation. G.) Conductance: Voltage curves plotted before (0 seconds) and after 1 second, 3 seconds, 8 seconds, 15 seconds exposure to U.V. irradiation and a dose dependent change is observed with the channels held at 0mV during U.V. irradiation. H.) Change in V0.5 after 15 seconds U.V. exposure (filled symbols:-100mV, empty symbols: 0mV holding potential). I.) Time to apparent half maximum current of voltage trace (-40mV opening potentials, (black filled: -100mV irradiation, black empty: 0mV irradiation).

**Figure 11:**
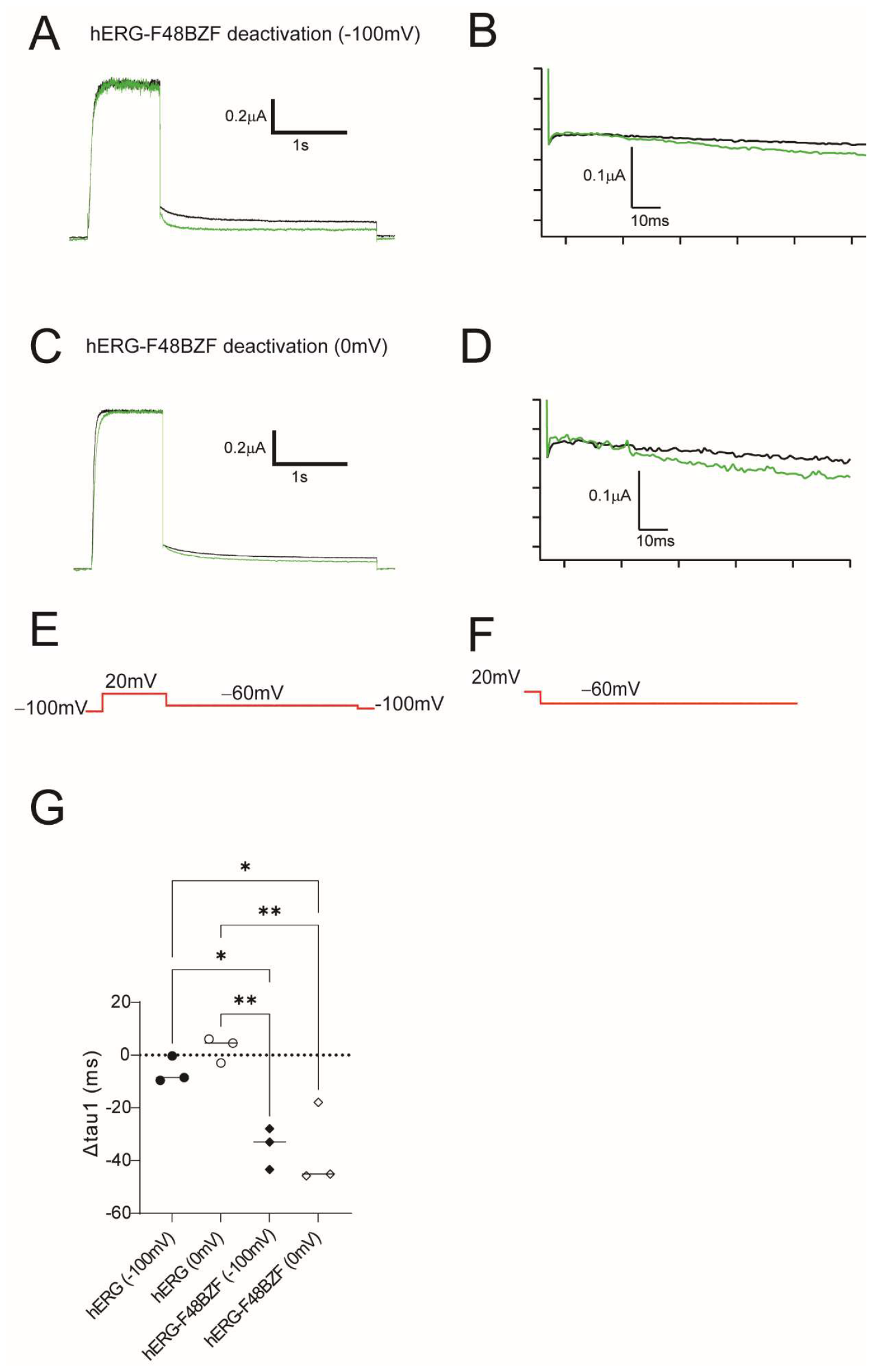
hERG-F48BZF channel biophysical properties of deactivation before and after U.V. irradiation. A.) Currents recorded from inside out excised patches containing hERG-F48 channels to measure rate of channel closure at -60mV. Channels held at -100mV, before U.V. exposure (black) and after 15 seconds U.V. irradiation (green), normalized. B.) The first 60ms of trace (zoomed in for clarity) of currents in panel A after channel closure at -60mV C.) Currents recorded from inside out excised patches containing hERG channels to measure rate of channel closure at -60mV. Channels held at 0mV, before U.V. exposure (black) and after 15 seconds U.V. irradiation (green), normalized. D.) The first 60ms of trace (zoomed in for clarity) after channel closure at -60mV of currents in panel C. E. Voltage protocol for panel A, C. F.) Voltage protocol for panels B, D. G.) Tau1 of deactivation of hERG-F48BZF channels before (black-filled) and after (black-empty) U.V. irradiation at respective holding potentials.

hERG-E50BZF photochemical crosslinking slows channel activation, speeds deactivation, and right shifts the voltage dependence of activation in a U.V dependent and state dependent manner (Fig.12,13). This suggests that the PAS domain motions altered here by crosslinking hERG-E50BZF are i) state dependent ii) play a role in channel activation at low voltages iii) alters both steady-state properties of the channel and kinetics of the channel.

**Figure 12:**
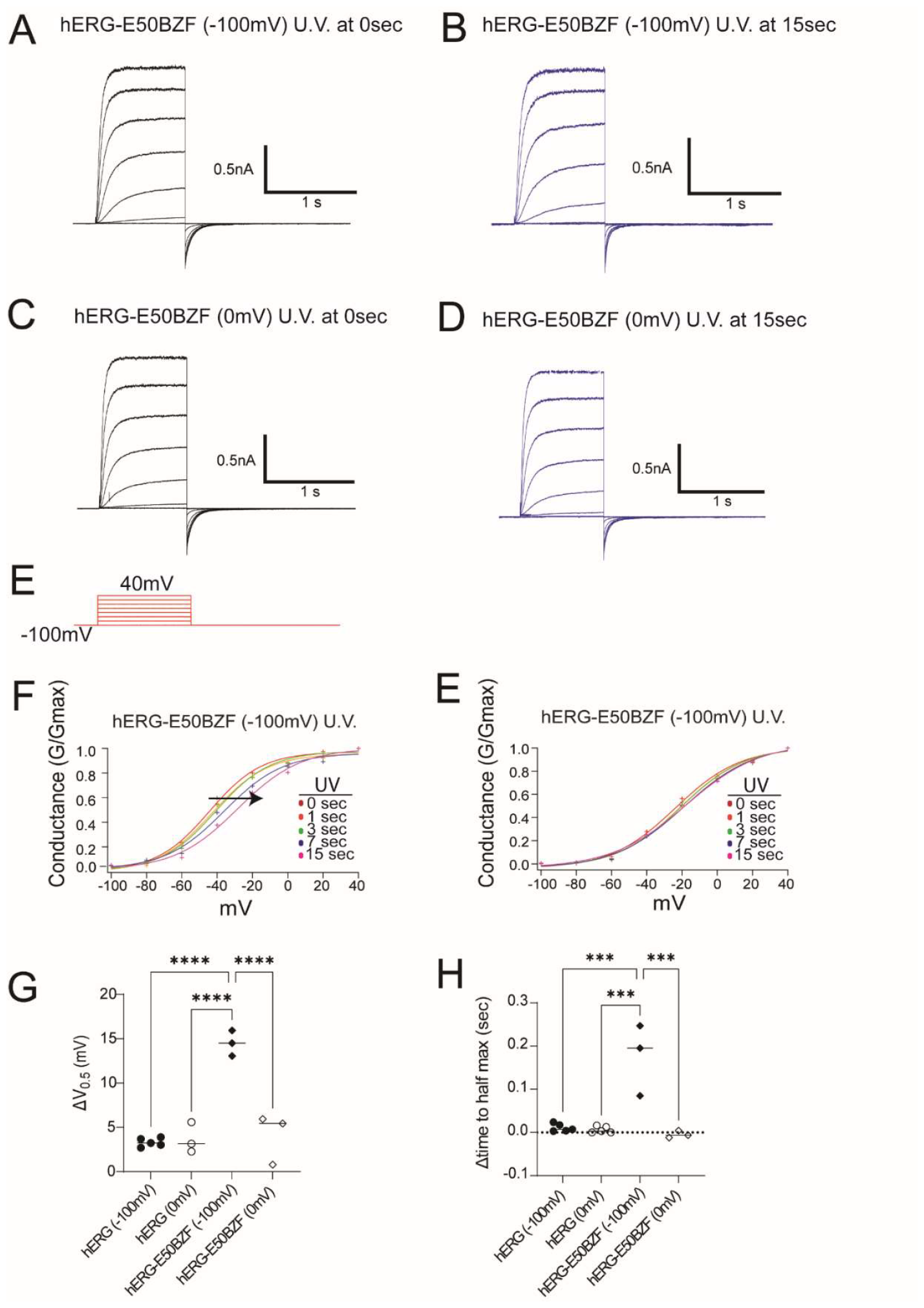
hERG-E50BZF channel biophysical properties of conductance and activation before and after U.V. irradiation. A.) Current family of traces recorded from inside out excised patches containing hERG-E50BZF channels held at -100mV prior, and during, U.V. exposure (U.V. 0 sec). B.) Family of traces as in panel A after 15sec cumulative exposure of U.V. irradiation. C.) Family of traces as in panel A with channels held open (0mV) and prior to U.V. irradiation D.) Family of traces as in panel C with channels held open (0mV) and after 15sec cumulative exposure to U.V. irradiation. E.) Voltage protocol for A-D. F.) Conductance Voltage curves plotted before (0 seconds) and after 1 second, 3 seconds, 8 seconds, 15 seconds exposure to U.V. irradiation and a dose dependent change is observed with the channels held at -100mV during U.V. irradiation. G.) Conductance: Voltage curves plotted before (0 seconds) and after 1 second, 3 seconds, 8 seconds, 15 seconds exposure to U.V. irradiation and no dose dependent change is observed with the channels held at 0mV during U.V. irradiation. H.) Change in V0.5 after 15 seconds U.V. irradiation (filled symbols:-100mV, empty symbols: 0mV holding potential). I.) Time to apparent half maximum current of voltage trace (-40mV opening potentials, (black filled: -100mV irradiation, black empty: 0mV irradiation).

**Figure 13:**
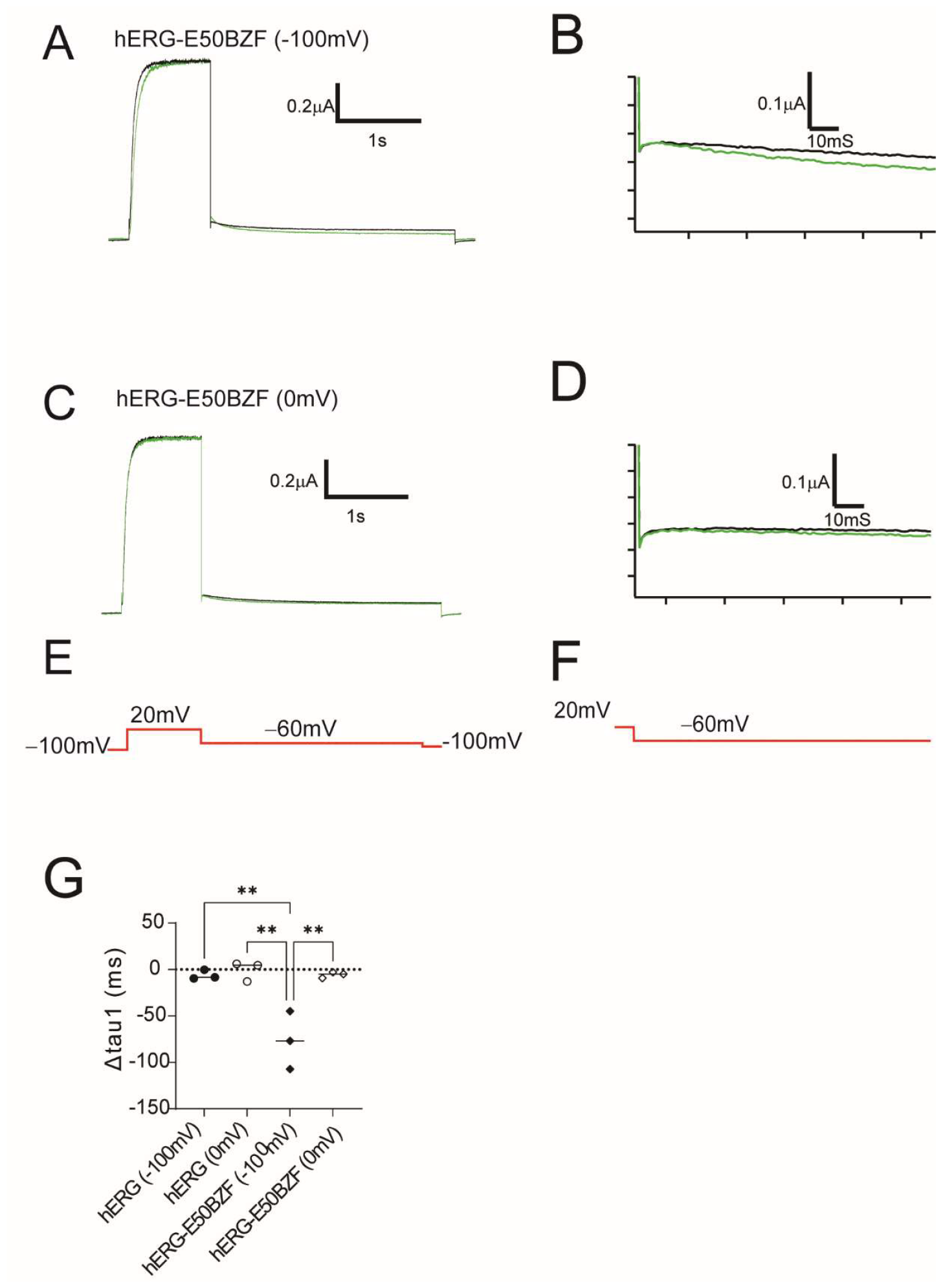
hERG-E50BZF channel biophysical properties of deactivation before and after U.V. irradiation. A.) Currents recorded from inside out excised patches containing hERG-E50 channels to measure rate of channel closure at -60mV. Channels held at -100mV, before U.V. exposure (black) and after 15 seconds U.V. irradiation (green), normalized. B.) The first 60ms of trace (zoomed in for clarity) of currents in panel A after channel closure at -60mV C.) Currents recorded from inside out excised patches containing hERG channels to measure rate of channel closure at -60mV. Channels held at 0mV, before U.V. exposure (black) and after 15 seconds U.V. irradiation (green), normalized. D.) The first 60ms of trace (zoomed in for clarity) after channel closure at -60mV of currents in panel C. E. Voltage protocol for panel A, C. F.) Voltage protocol for panels B, D. G.) Tau1 of deactivation of hERG-E50BZF channels before (black-filled) and after (black-empty) U.V. irradiation at respective holding potentials.

How these hERG-BZF crosslinks alter the conformational landscape of PAS domain remains to be shown. The cryo-EM structures of hERG and the co-crystal of the PAS-CNBHD indicate that residues in the PAS (E50) and CNBHD domain (K741) are within structural proximity to form crosslinks (Fig.#) (22). But the extent of which these residues interact during gating remains to be determined. It is also possible that photo reactive crosslinking could occur intra domain (within the PAS domain itself). Regardless, that the PAS-CNBHD interaction has been shown to be integral to hERG deactivation gating, and that both the structural interactions and functional regulation of the channel via this interface are sensitive to mutation, suggests that this interaction is crucial and tunes hERG channels to its physiological role in the heart (13, 23). It now appears that the effect of mutations in the PAS domain could be attributed to the altered conformational landscape of the PAS domain. Future studies may determine whether the crosslinking effect is intra-or inter-subunit and modeling may provide insight on the structure-function effects observed in this study.

The approach utilized a genetically encoded non-canonical amino acid (ncAA) as small and silent probe to study ion channel dynamics in native membranes under voltage control to simultaneously record ion channel energetic and structural information (24). With photo crosslinking we tested if the conformation of the PAS domain in hERG channels allosterically regulated the channel gate. Taken together, the results observed here indicate that photo cross-linking channels with BZF in the PAS domain destabilized the open state of the channel and stabilized the closed state of the channel. Crosslinked hERG-BZF channels were slower to activate, faster to deactivate and opened at more positive voltages (Table 1). We hypothesize that the PAS domain motions are critical to the rate of channel activation, deactivation, and allosterically affects the rate and voltage at which the channel gate opens *and* closes (Fig.14). While it was previously known that the PAS domain can regulate the rate to which the channel closes, here we learn that the rate of channel closure is dependent on the conformation of the PAS domain. Additionally, our results suggest that the PAS domain motions couple the voltage sensor to the pore.

**Table 1:**
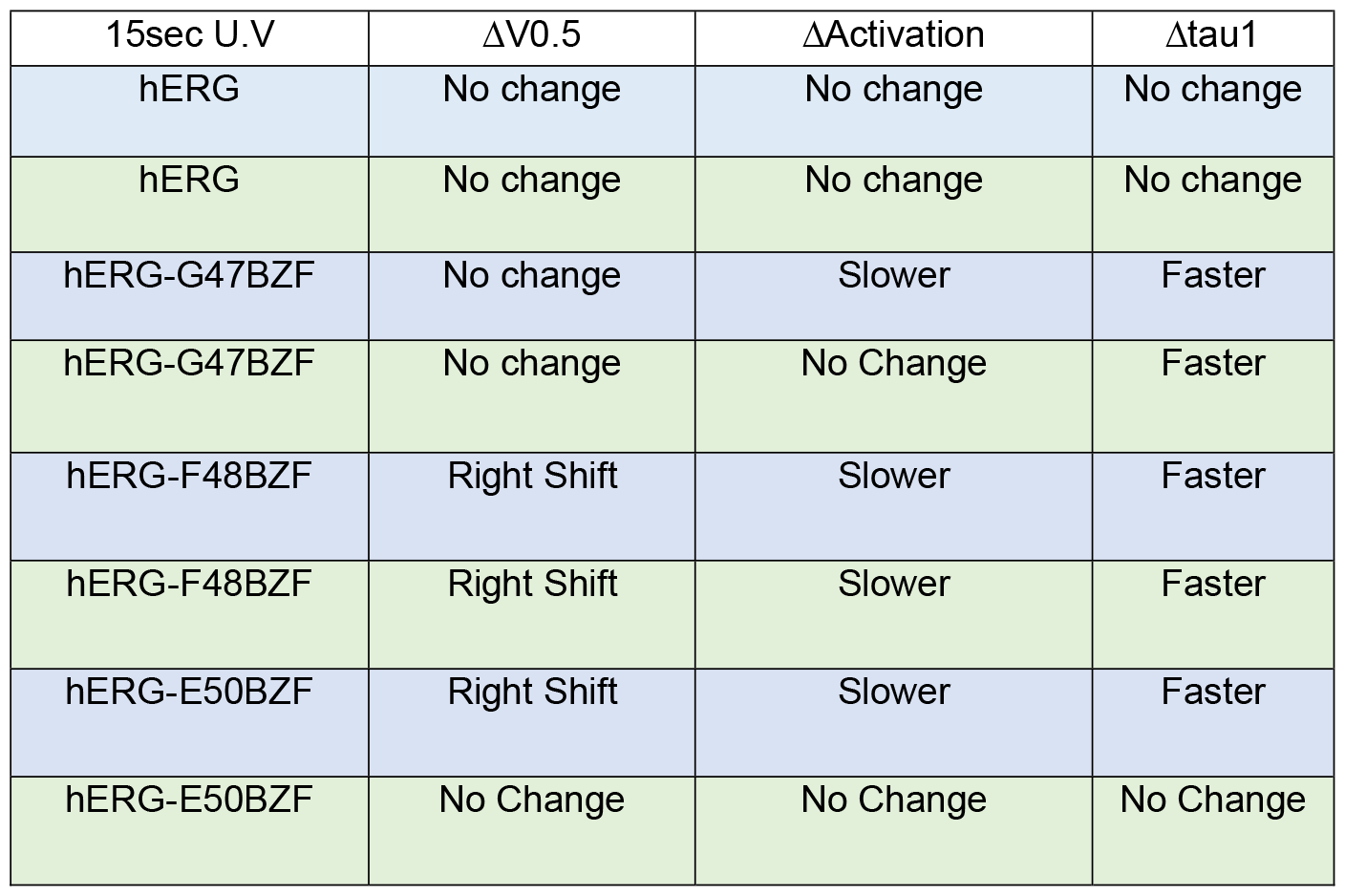
Summary of Changes in response to U.V. irradiation of hERG-BZF channels (Blue -100mV holding voltage, Green 0mV holding voltage. Results summarized from crosslinking studies.

| 15sec U.V | $\Delta V_{0.5}$ | $\Delta \text{Activation}$ | $\Delta \tau_{1}$ |
| --- | --- | --- | --- |
| hERG | No change | No change | No change |
| hERG | No change | No change | No change |
| hERG-G47BZF | No change | Slower | Faster |
| hERG-G47BZF | No change | No Change | Faster |
| hERG-F48BZF | Right Shift | Slower | Faster |
| hERG-F48BZF | Right Shift | Slower | Faster |
| hERG-E50BZF | Right Shift | Slower | Faster |
| hERG-E50BZF | No Change | No Change | No Change |

**Figure 14:**
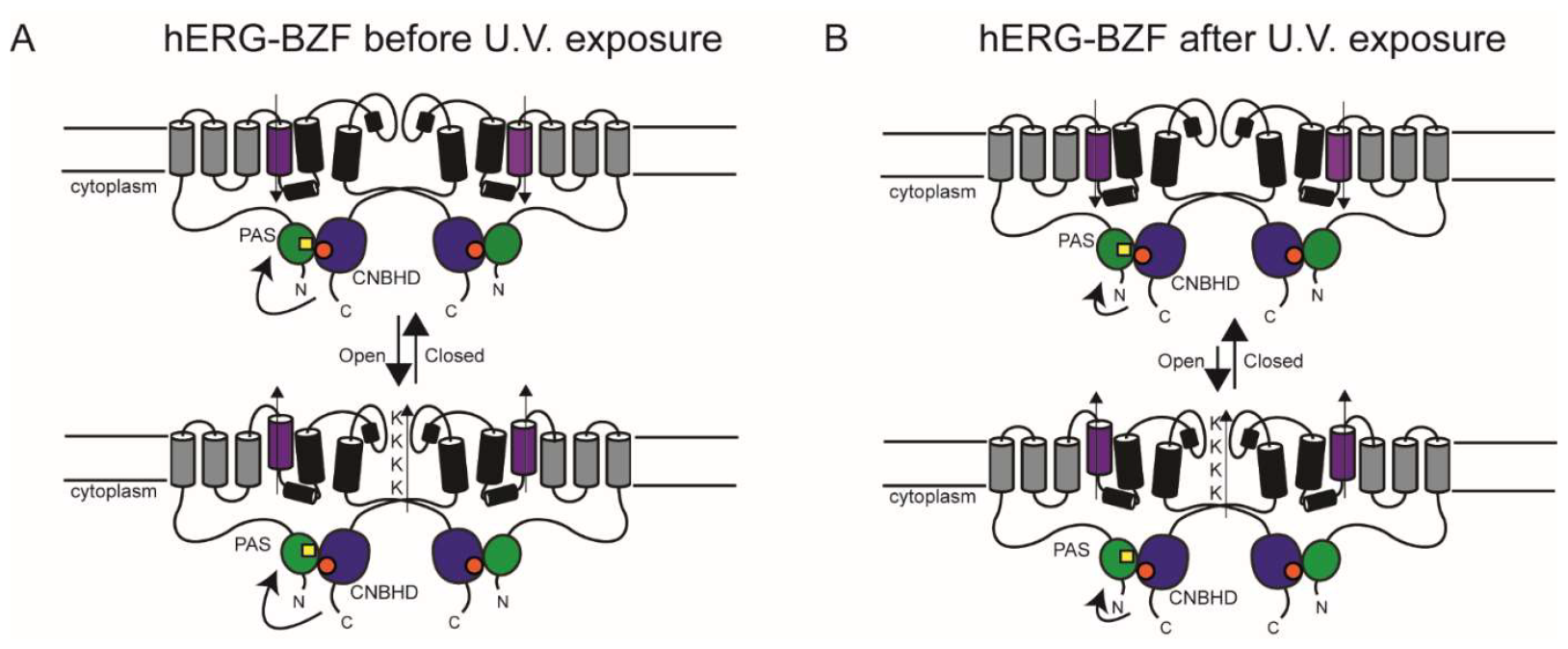
hERG schematic color coded as in Figure 1, BZF depicted by yellow square in the PAS domain. A.) Prior to U.V. irradiation PAS domain with BZF incorporated can regulate the channel via dynamic motions native to the domain. B.) After U.V. irradiation the PAS domain with BZF incorporated can form crosslinks (depending on location of incorporation or irradiation state, see Table 1), that reduce PAS domain motions and stabilize the closed state of the channel.

## Materials and Methods

### Molecular Biology

Full length human ERG construct (protein Accession NP_000229) expression clone with inactivation removed (S620T variant) was subcloned into a modified pcDNA3 vector that contained a T7 promoter and 3’ and 5’ untranslated region of a Xenopus B-globin gene, protein is denoted hERG in this manuscript. Point mutations to insert TAG codons into the hERG gene were generated with site directed mutagenesis by BioInnovatise, Rockville MD and validated by Sanger sequencing. All hERG and hERG-BZF constructs in this manuscript had a C-terminal Citrine fusion tag. The orthogonal set encoding the BZF-tRNA and BZF RNA synthetase (RS) were a generous gift from Thomas P. Sakmar at The Rockefeller University. The channel RNA was synthesized in vitro using mMESSAGE mMACHINE T7 transcription kit (ThermoFisher, Waltham, MA) from the linearized cDNA.

### Oocyte Harvest, Injection, and Incubation

Oocytes were harvested from Xenopus *laevis* as previously described (Varnum et al. 1995) or purchased from Ecocyte Biosciences. Channel mRNA was injected into the cytosolic region of the oocyte (∼50nL). The BZF plasmids that encoded the orthogonal tRNA and aminoacyl-tRNA synthetase specific to TAG codon suppression and incorporation of BZF were blind injected into the oocyte nucleus (10nL at a ratio of 40ng/uL RS:120ng/uL tRNA). Oocytes were incubated at 16ºC in ND96 with 50μg/mL gentamycin and 2.5mM sodium pyruvate for 2-4 days and rocked with gentle agitation. BZF (Apollo Scientific or BaChem) was dissolved in 1M NaOH and added to the bath at 1mM (pH adjusted to 7.6 with 1N HCl).

### Channel Expression Screening

To validate hERG-G47TAG, hERG-F48TAG, hERG-E50TAG (hERG-TAG) channels for TAG codon suppression and BZF incorporation, channels were allowed to express in the presence *or* absence of BZF. Using two-electrode-voltage-clamp, hERG-TAG currents were recorded from oocytes. hERG-TAG channels that expressed in the presence of BZF and with robust current density at 2DPI-4DPI, but not the absence of BZF, were used for these experiments.

### Electrophysiology and Photo crosslinking

Excised inside out patch clamp recordings from oocytes expressing hERG and hERG-BZF channels were performed 3-4 days after injection using a Nikon Ti2 microscope with a 40X water immersion objective (N.A. 1.15). BZF was excited with wide-field excitation using a Nikon Elements controlled SOLA SM II and a chroma 460/50m excitation filter. Excised patch membranes were exposed to U.V. light at discrete time intervals and holding voltages (-100mV or 0mV). After each discrete time interval of U.V. exposure biophysical recordings were performed. The bath solution contained 130mM KCl, 0.2M EDTA, 10mM HEPES, pH 7.4. The internal solution contained KCl 4mM, NaCl 126mM, EDTA 0.2mM and HEPES 10mM, pH 7.2. The initial pipette resistance was 0.5-0.9MOhms for oocyte recordings. Borosilicate patch electrodes were made using a P97 micropipette puller (Sutter Instrument, Novato, CA) and polished with a Narishige MF2 microforge. Recording were made at 22ºC to 24ºC.

The channel conductance-voltage relationship (G-V curve) was measured from tail currents at -100mV. It was fitted with a sigmoid function in Igor Pro and the change in voltage at 50% conductance is reported. This was calculated by taking the difference between V0.5 (15 seconds irradiation)-V0.5 (0 seconds irradiation)=ΔV0.5.

The channel activation was measured as apparent time to half max current of the trace invoked by a -40mV opening potential. The difference in apparent time to half max current was calculated by taking the difference in values at 15 seconds and 0 seconds irradiation at respective holding potentials during irradiation.

The channel deactivation rate was measured from current relaxations with repolarizing voltage steps (-60mV in this study) and fit with an exponential function (y=Ae^(-t/tau)) where t is time and tau is the time constant. The change in tau1, Δtau1=tau1 (15 seconds irradiation)-tau1 (0sec irradiation).

Changes reported in measured biophysical values are changes in the mean differences between 15 seconds U.V. irradiation and 0 seconds U.V. irradiation to highlight U.V. dependent (or independent) changes in channel properties due to BZF crosslinking.

Data was plotted in Igor Pro or Graphpad.

## Statistics

Statistics reported here are calculated from student’s T test or one-way anova where appropriate. Asterisk (P<=0.05, *), (P<=0.01, **), (p<0.001, ***), plots were generated in Graph Pad.

## Acknowledgements

We thank Dr. Ivy Dick, Dr. W.N. Zagotta, Dr. Ashley A. Johnson, and Dr. M. Bamgboye for their helpful discussions. We expressly thank Dr. Sharona E. Gordon for her keen expertise in editing and scientific knowledge in channel expression with ncAA, her support was unwavering, sharp, and critical to the success of this manuscript. This work was supported by NIH Grant R01-130701-01 and R01-12752301A1 (M.C.T.) and training grant K99GM144684 (S.J.C.).

The authors declare no competing financial interests.

## Author contributions

S.J. Codding collected, analyzed, designed the experiments and wrote the paper. M.T. Trudeau designed the initial experiments for this manuscript and co-wrote the paper.

